# Supracategorical fear information revealed by aversively conditioning multiple categories

**DOI:** 10.1101/2020.05.21.107805

**Authors:** Seth M. Levine, Miriam Kumpf, Rainer Rupprecht, Jens V. Schwarzbach

**Affiliations:** Department of Cognitive and Clinical Neuroscience, Central Institute of Mental Health, Medical Faculty Mannheim, Heidelberg University, 68163 Mannheim, Germany; Department of Psychiatry and Psychotherapy, University of Regensburg, 93053 Regensburg, Germany

**Author notes:** Equal contribution (Seth M. Levine, Miriam Kumpf).

**Keywords:** aversive-learning, fear-conditioning, fMRI, multivariate pattern analysis, representational similarity analysis

## Abstract

Fear-generalization is a critical function for survival, in which an organism extracts information from a specific instantiation of a threat (e.g., the western diamondback rattlesnake in my front yard on Sunday) and learns to fear—and accordingly respond to—pertinent higher-order information (e.g., snakes live in my yard). Previous work investigating fear-conditioning in humans has used functional magnetic resonance imaging (fMRI) to demonstrate that activity-patterns of stimuli from an aversively-conditioned category (CS+) are more similar to each other than those of a neutral category (CS-). Here we designed a three-phase (i.e., baseline, conditioned, extinction) experiment using fMRI and multiple aversively-conditioned categories to ask whether we would find only similarity increases within the CS+ categories or also an increase in similarity between the CS+ categories. Using representational similarity analysis, we correlated a set of models to activity-patterns underlying several regions of interest and found that, following fear-conditioning, between-category and within-category similarity increased for the CS+ categories in the superior frontal gyrus (SFG) and the right temporal pole (rTP). Activity patterns in the object-selective lateral occipital cortex tended to prefer the semantic model, regardless of the experimental phase. These results advance prior pattern-based neuroimaging work by exploring the effect of aversively-conditioning multiple categories and indicate an extended role for the SFG and rTP in potentially linking discrete information or abstractly representing supracategorical information during fear-learning for the purpose of proper generalization.

## Introduction

Fear-learning is an essential process for an organism’s survival, as it allows for a reasonable response to environmental threats. Classically, this process has been studied through fear-conditioning, a form of Pavlovian conditioning, in which an organism is exposed to a neutral stimulus (e.g. a picture of a dandelion) that is paired with an aversive, unconditioned stimulus (US, e.g., a mildly-painful electric shock); following a sufficient number of such pairings, the neutral stimulus is rendered a conditioned stimulus (CS+) and, even absent the US, yields the same physiological fear-response in the organism [1, 2].

This basic process gives rise to the higher-order process of fear-generalization. Recognizing that an entity in the environment might be dangerous based on its similarity to other previously-encountered entities is a critical extension of fear-learning. With respect to the previous example, fear-generalization could result in a fear-response upon seeing other plants besides dandelions. While the physiological process of fear-generalization may prove helpful in some situations, it stands to reason that a faulty generalization process could explain a variety of psychiatric disorders that are characterized by elevated levels of fear or anxiety [3, 4]. As anxiety disorders have a lifetime prevalence of 16.6% [5], investigating the neural mechanisms of fear-generalization in humans is of particular interest to the translational neuroscience community.

Numerous neuroimaging studies have demonstrated that the amplitude of the blood-oxygen-level-dependent (BOLD) signal evoked by a CS+ differs from that evoked by a neutral stimulus (CS-) in various regions of the human brain (see [6] for a review). In terms of multivariate pattern analysis (MVPA) [7], recent work has shown increased similarity between activity patterns evoked by a CS+ and those evoked by images of spiders in the right anterior temporal lobe of humans with arachnophobia [8]. Other studies employing pattern-based analyses have shown that the activity patterns evoked by a CS+ in the insular cortex bear a closer resemblance to those evoked by the US [9] or that the activity patterns representing stimuli belonging to the same semantic category (e.g., animals or tools) in object-selective cortex are more similar to each other after some elements of the category have been paired with an electric shock [10]. Following especially from this last example, one could surmise that, given the increased within-category similarity, the human brain has the cognitive ability to generalize a specific, conditioned fear to a superordinate group. We were interested in building on this finding and investigating whether fear-conditioning could also elicit increased pattern similarity between semantically *unrelated* categories.

To this end, we conducted a three-phase experiment (i.e., baseline, conditioned, extinction) in which participants viewed images from three categories (i.e., animals, produce, and tools) while lying in the MR scanner. During the conditioned phase, 50% of the images from two of the categories were paired with mildly aversive electric shocks, while the third category remained neutral throughout the entire experiment. Within the framework of representational similarity analysis (RSA) [11], we used a set of models to explain how the pattern similarity of the stimuli might change as a function of fear-conditioning. Specifically, we were interested in whether we would only see increases in similarity *within* the CS+ categories or (also) *between* the CS+ categories. The models make different assumptions about the underlying operating principles of fear-generalization, with respect to categorical knowledge, and can therefore reveal differences in information processing of distinct brain regions that are involved in learning, affect, and semantics.

## Methods

### Participants

Twenty-one participants (6 males, 15 females; mean age = 25.2 yrs, age range = 20 – 30 yrs) were recruited from the local community. Participants had no current neuropsychiatric diagnoses, were not taking any psychotropic medication, and provided written informed consent before taking part in the study. All experimental procedures were approved by the ethics committee of the University of Regensburg and complied with the Declaration of Helsinki.

### Stimuli

Stimuli used in the experiment comprised three distinct categories: animals, produce (i.e., fruits/vegetables), and tools. Thirty unique images from the three categories were obtained from multiple sources, either contributed by other researchers [12] or freely available online [13, 14] (www.pexels.com, www.unsplash.com). We avoided using potentially threatening images (e.g., knives/spiders) to lessen arousal-based effects.

The visual stimuli were presented with A Simple Framework [15], built on the Psychophysics toolbox [16] and MATLAB R2015b (Mathworks, Natick, USA), and cast onto a semitransparent screen behind participants using an LED projector (PROPixx, VPixx Technologies Inc., Saint Bruno, Canada) at a frame rate of 60 Hz and a resolution of 1024 × 768 pixels.

Electrical stimuli (duration = 2 ms) were delivered via a DS7A current stimulator (Digitimer Limited, Letchworth Garden City, UK), the timing of which was controlled by ASF and an Arduino^®^ (Arduino SA, Chiasso, Switzerland).

### Experimental design

#### Pre-scan protocols

Participants first completed the German version of the State-Trait Anxiety Inventory [17], which are two questionnaires each containing 20 self-reported items used to assess state and trait anxiety on a 4-point Likert scale.

Next, participants completed a threshold-acquisition session, in which they applied electric shocks to their left wrists, while increasing the amperage, until they found a near-painful sensation that they considered tolerable but unpleasant. The individualized amperages (mean = 11.13 mA, range = 1.5 – 29.0 mA) obtained from this session were used during the conditioned phase of the main experiment.

#### Main experiment

The main experiment contained three phases: baseline, conditioned, and extinction. Each phase followed the same event-related protocol, with the exception that participants received electric shocks during the conditioned phase. During a given phase, participants viewed 90 stimuli in total (i.e., 30 stimuli from each category). To prevent discomfort, rather than having participants undergo one long run (per phase) with 90 stimulus presentations, each phase was split into three shorter runs, in which we presented 30 stimuli (i.e., 10 stimuli per category) per run. A given run thus contained 30 trials; each trial was 3 s in duration (2 s of stimulus presentation + 1 s of fixation), followed by a temporally-jittered intertrial interval of 6 + X s, with X ∼ geom(0.3), truncated at 10 s, yielding possible intertrial intervals of 6 to 10 s in steps of 0.5 s. Each run started with a 12 s fixation period, to account for T1 saturation, and ended with a 12 s fixation period to capture the BOLD response from the final trial. In total, a given run lasted between 4 min 48 s and 6 min 44 s, depending on the intertrial jitter, with an average run length of approximately 5 min 25 s. A black fixation dot was always present in the center of the screen.

During all runs, to ensure that participants were paying attention to the visual stimuli, we asked them to perform a one-back task, in which they indicated via button-press whether the image they viewed on a given trial was from the same category (right index finger response) or a different category (right middle finger response) as the image from the previous trial. Participants’ responses were recorded starting from the onset of the stimulus presentation to the end of the 1 s fixation period that followed the stimulus.

Additionally, during the conditioned phase, electrical stimulation was administered at the offset of stimulus presentation for 50% of images that belonged to two of the three categories. In each run of the conditioned phase, there were thus 10 electrical shocks (i.e., 5 shocks per conditioned category); the first presentations of exemplars from conditioned categories co-terminated with electric shocks, to ensure that the conditioning was sufficiently re-instated between runs. The remaining four shocks per category (per run) were pseudorandomized across the remaining presentations of the stimuli of those categories. Of the 21 participants, seven were conditioned to animals and produce, seven were conditioned to animals and tools, and seven were conditioned to produce and tools.

### Neuroimaging data acquisition

The neuroimaging experiment was carried out at the University of Regensburg, Germany using the 3T research-dedicated MR scanner (Magnetom Prisma, Siemens, Erlangen, Germany) and a 64-channel head coil. Functional images were acquired with a T2*-weighted EPI sequence (64 slices per volume, acquired with a multiband factor of 4 [18, 19], field of view (FOV) = 192 × 192 mm2, isotropic voxel resolution (VR) of 2 × 2 × 2 mm3, no interslice gap, repetition time (TR) = 2000 ms, echo time (TE) = 30 ms, flip angle (FA) = 75°). Four dummy scans from the beginning of each functional run were acquired to account for signal saturation. For a given run of the main experiment, we acquired between 144 to 202 volumes, depending on the intertrial jitter (see section Main experiment).

For co-registration of the functional images to high-resolution anatomical images, we acquired 160 slices of a T1-weighted magnetization-prepared rapid gradient-echo (MPRAGE) sequence (FOV = 256 × 256 mm2, VR = 0.9766 × 0.9766 × 1 mm3, TR = 1910 ms, TE = 3.67 ms, FA = 9°) for each participant.

### Data analysis

Neuroimaging data were analyzed with the FMRIB Software Library (FSL) [20] and the CoSMoMVPA toolbox [21] for MATLAB.

#### Preprocessing

Preprocessing of the functional data included removal of non-brain tissue [22], slice time correction, motion correction with respect to the middle volume of each run (using 6 degrees of freedom and trilinear interpolation), and high-pass filtering (cutoff = 100 s). No spatial smoothing was performed. We co-registered each participant’s functional data to the corresponding high-resolution structural scan using 7 degrees of freedom [23] and then to standard space (i.e., MNI152 2mm) using 12 degrees of freedom.

#### First-level modeling

For each run of the experiment, time-course data from each voxel were analyzed using the general linear model (GLM) containing 30 regressors of interest (i.e., one regressor per image presented to the participant) and six motion correction parameters (i.e., 3 translations, 3 rotations) modeled as regressors of non-interest. Hemodynamic response functions were generated by convolving the regressors of interest with gamma functions using FSL’s default parameters *ϕ* = 0 s, *σ* = 3 s, mean lag = 6 s). The resulting t-scores of the beta weights per regressor were concatenated across the three runs of an experimental phase, resulting in 90 whole-brain t-score maps (per phase), reflecting the 90 stimuli that a participant viewed.

#### Region-of-interest definition

Regions-of-interest (ROIs) comprised the amygdala, posterior fusiform gyrus, insula, inferior lateral occipital cortex (LOC), superior frontal gyrus (SFG), and right temporal pole (rTP), which we selected based on previous neuroimaging studies that similarly combined forms of MVPA [7] with aversive-learning [8–10, 24]. The ROIs were defined using the Harvard-Oxford probabilistic (sub)cortical atlas [25, 26] and contained voxels that belonged to the given region with a probability greater than 0.5 (see Fig. 1).

**Fig. 1.**
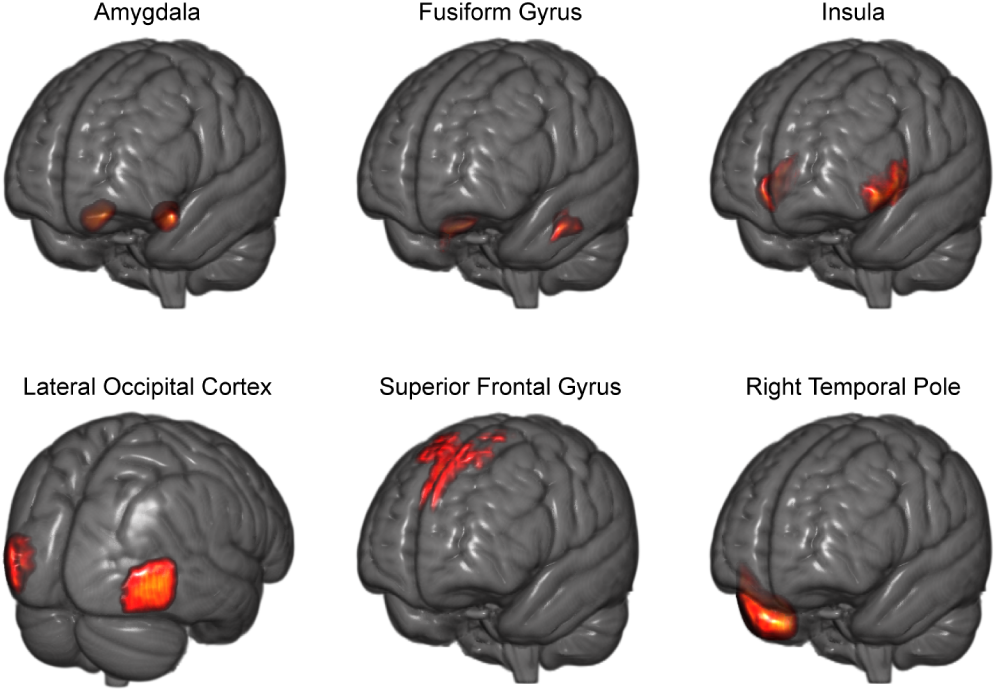
Regions of interest (ROIs) used in the current study based on previous fear-learning investigations that demonstrated conditioning-based modulations of multivoxel patterns. See section Region-of-interest definition for details about the ROI selection and definition.

#### Representational similarity analysis

Within the six ROIs, we were mainly interested in determining whether, following fear-conditioning, activity-patterns would become more similar between conditioned categories or only within conditioned categories. To this end, we constructed five 90 × 90 model dissimilarity matrices (DSMs) that each contained zeros (indicating similarity between two stimuli) and ones (indicating dissimilarity between two stimuli). The five model DSMs reflected the following stimulus relationships: between + within CS+, between CS+, within CS+, within CS-, and semantic similarity (the last two models served as controls). The DSMs involving conditioned stimuli were participant-specific (as different participants were conditioned to different categories); the semantic DSM was the same for all participants (see Fig. 2).

**Fig. 2.**
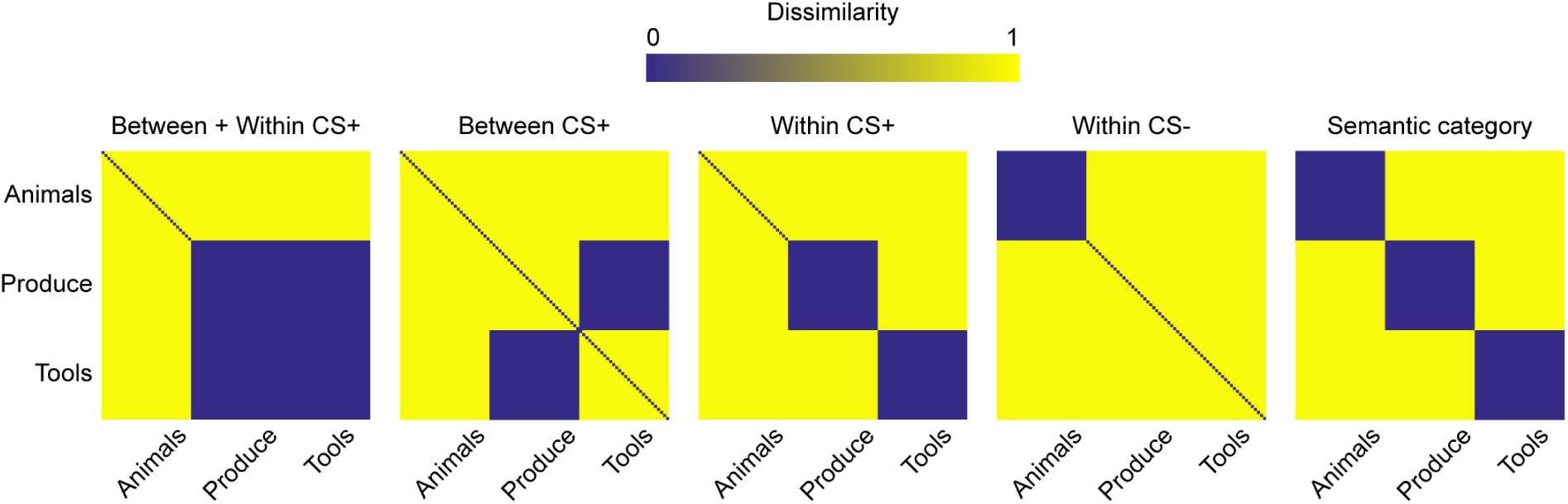
A visual depiction of the five model dissimilarity matrices (DSMs) used in the representational similarity analysis. Each 90 × 90 matrix reflects the binarized dissimilarity between each pair of the 90 stimuli that a participant viewed during a given phase of the experiment. Together the five DSMs represent distinct hypotheses regarding the relationship among the stimuli from three distinct semantic categories (i.e., 30 animals, 30 fruits/vegetables, and 30 tools), which we used to test whether aversive-learning would yield increases in within-category or between-category similarity (or both) at each of the six ROIs. As the models were specific to certain participants (depending on which categories were fear-conditioned), depicted here are example DSMs from a participant for whom produce and tools served as the two CS+ categories. One-third of the participants were conditioned to animals and produce, one-third to animals and tools, and one-third to produce and tools. For all participants, the rightmost DSM (semantic category) was identical and served as a control model.

For a given participant, we computed the pairwise similarity of the 90 activity patterns within each ROI (per phase; all 270 patterns for a given ROI were mean-centered) using (Pearson) correlation distance (i.e., 1 – r), which yielded a neural DSM reflecting the underlying representational space of these 90 stimuli in the given ROI. We then Spearman correlated our five model DSMs with the neural DSMs of the six ROIs, for each phase. This procedure was carried out for all participants, and the resulting 90 correlations (per participant) were Fisher-transformed and submitted to a three-way repeated-measures analysis of variance (ANOVA) containing factors ROI (6 levels) × Phase (3 levels) × Model (5 levels) computed with SPSS ver. 25 (IBM Corp., Armonk, NY, United States). Ultimately, we wanted to determine whether there was a three-way interaction between these factors, which would indicate that different model DSMs explain activation patterns in different experimental phases for different ROIs. Additionally, because we were specifically interested in determining whether representational spaces were modulated as a function of fear-conditioning, we carried out paired t-tests for each model, testing whether the resulting correlation differed between the conditioned phase and the baseline phase. Computing these five t-tests at each of the six ROIs yielded 30 statistical tests, which we accounted for with a False Discovery Rate (FDR) correction [27]).

#### Anxiety correlations

Lastly, we wanted to determine whether there was a relationship between the change in the explanatory value of the RSA models identified from the FDR-correction and the participants’ STAI scores. To this end, we carried out a set of post-hoc correlations between the STAI results and the correlation changes (from baseline to conditioned), of the Bw+Wi, Bw, Wi, and CS-models in the SFG and the Bw+Wi model in the rTP.

## Results

### Increase in similarity both within and between conditioned categories in the superior frontal gyrus and temporal pole

In order to initially discern whether there were overall differences between the models, the 90 correlation values that we obtained from correlating the activity patterns from six regions with the five models at three different phases were Fisher transformed and submitted to a three-way repeated-measures ANOVA. This procedure supported the observation of general differences among regions (F_5,100_ = 91.9, p = 8.42 × 10^−36^), phases (F_2,40_ = 4.87 p = 0.013), and models (F_4,80_ = 10.6, p = 6.08 × 10^−7^), in addition to regions yielding different correlations depending on the phase (Region × Phase: F_10,200_ = 3.96, p = 6.03 × 10^−5^), models performing differently depending on the region (Region × Model: F_20,400_ = 29.9, p = 6.96 × 10^−67^), and models performing differently depending on the phase (Phase × Model: F_8,160_ = 6.02, p = 9.30 × 10^−7^). Most critically, the ANOVA also supported the observed three-way interaction in that different models performed differently at different experimental phases, depending on the region (Region × Phase × Model: F_40,800_ = 1.98, p = 3.65 × 10^−4^). See Figure 3 and Table 1 for an overview of the results.

**Table 1.** Results from the three-factorial repeated-measures ANOVA

|  | F-score | df1, df2 | p-value |
| --- | --- | --- | --- |
| Region | 91.9 | 5, 100 | $8.42 \times 10^{-36}$ |
| Phase | 4.87 | 2, 40 | 0.013 |
| Model | 10.6 | 4, 80 | $6.08 \times 10^{-7}$ |
| Region × Phase | 3.96 | 10, 200 | $6.03 \times 10^{-5}$ |
| Region × Model | 29.9 | 20, 400 | $6.96 \times 10^{-67}$ |
| Phase × Model | 6.02 | 8, 160 | $9.30 \times 10^{-7}$ |
| Region × Phase × Model | 1.98 | 40, 800 | $3.65 \times 10^{-4}$ |

**Fig. 3.**
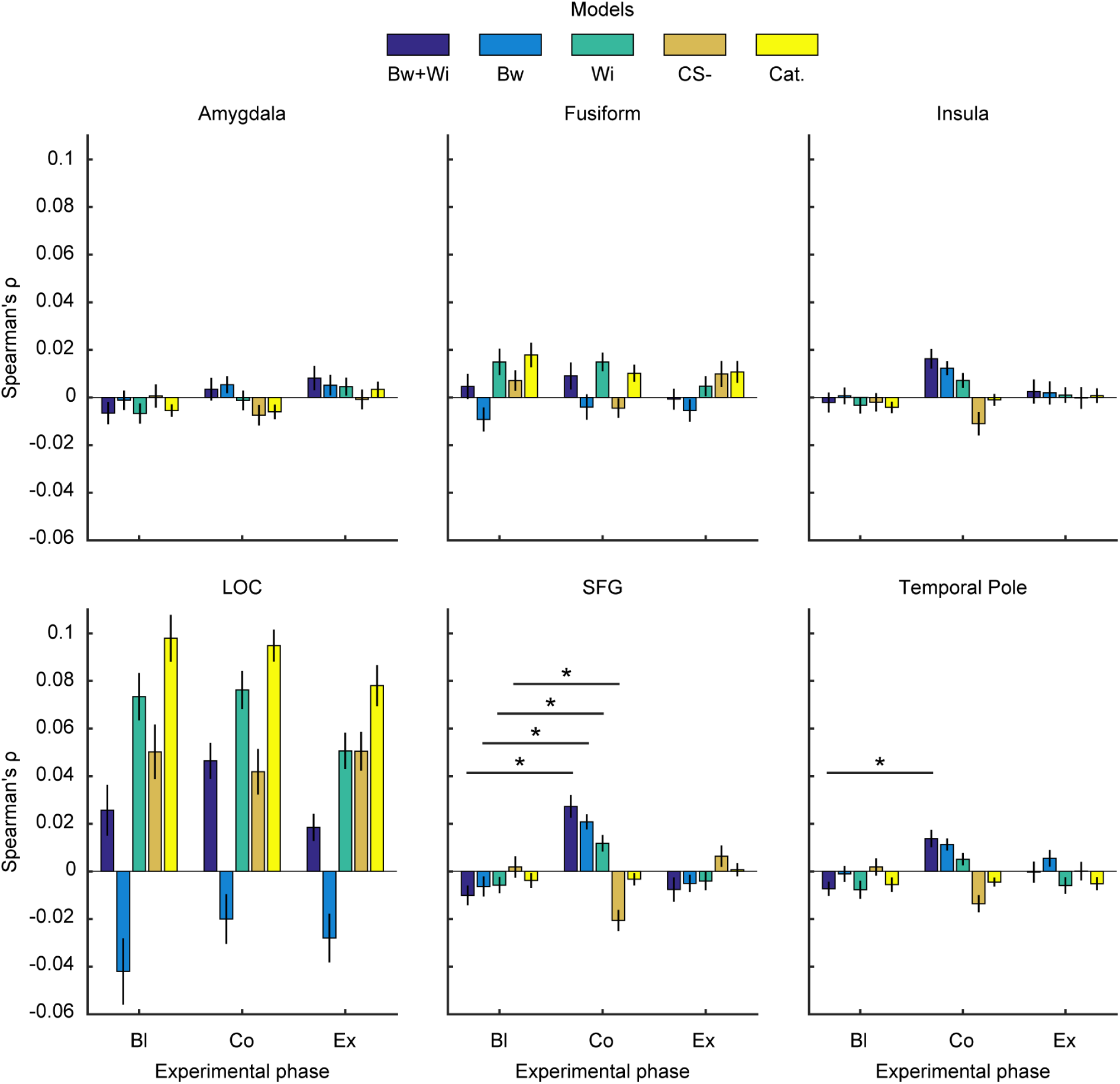
Spearman correlations of the activity patterns from each ROI (Fig. 1) with the five model DSMs (Fig. 2) at the three phases of the experiment. The resulting correlations exhibited a three-way interaction in that different models explained activity patterns of different ROIs differently in different experimental phases (as supported by a three-factorial repeated-measures ANOVA). FDR correction revealed that both between-and within-category (Bw+Wi) similarity increases best explained changing activity patterns in the superior frontal gyrus (SFG) and right temporal pole (rTP). The SFG also appeared to show increases in between-category similarity (Bw) and within-category similarity (Wi), and a decrease in within-category similarity of the CS-category. Interestingly, in all experimental phases, the lateral occipital cortex (LOC) preferred the semantic category model, and we did not find evidence for conditioning-based modulation within object-selective cortex. Error bars represent SEM.

As we were specifically interested in whether the performance of our models was modulated by fear-conditioning, we carried out a series of tests on the difference between each model’s correlation to a region’s activity patterns during the conditioned phase compared to the baseline phase. This procedure revealed that, following fear-conditioning, the Between+Within (Bw+Wi) CS+ model better explained the activity patterns within the superior frontal gyrus (SFG: t_20_ = 6.08, p = 6.06 × 10^−6^) and the right temporal pole (rTP: t_20_ = 4.00, p = 6.99 × 10^−4^). The only other models whose corresponding tests surpassed the FDR-corrected threshold of 0.0042 were the Between (Bw) CS+ model, the Within (Wi) CS+ model, and the Within CS-(CS-) model within respect to activity patterns in the SFG (t_20_ = 5.30, p = 3.50 × 10^−5^; t_20_ = 3.49, p = 0.0023; and t_20_ = −3.53, p = 0.0021, respectively). Although the overall pattern of results for the Insula were qualitatively similar to those of the SFG and rTP, the change in performance of the Bw+Wi model at the Insula did not surpass the FDR-corrected threshold (t_20_ = 2.71, p = 0.013). See Figure 3 for an overview and Table 2 for a full list of the corresponding results.

**Table 2.**
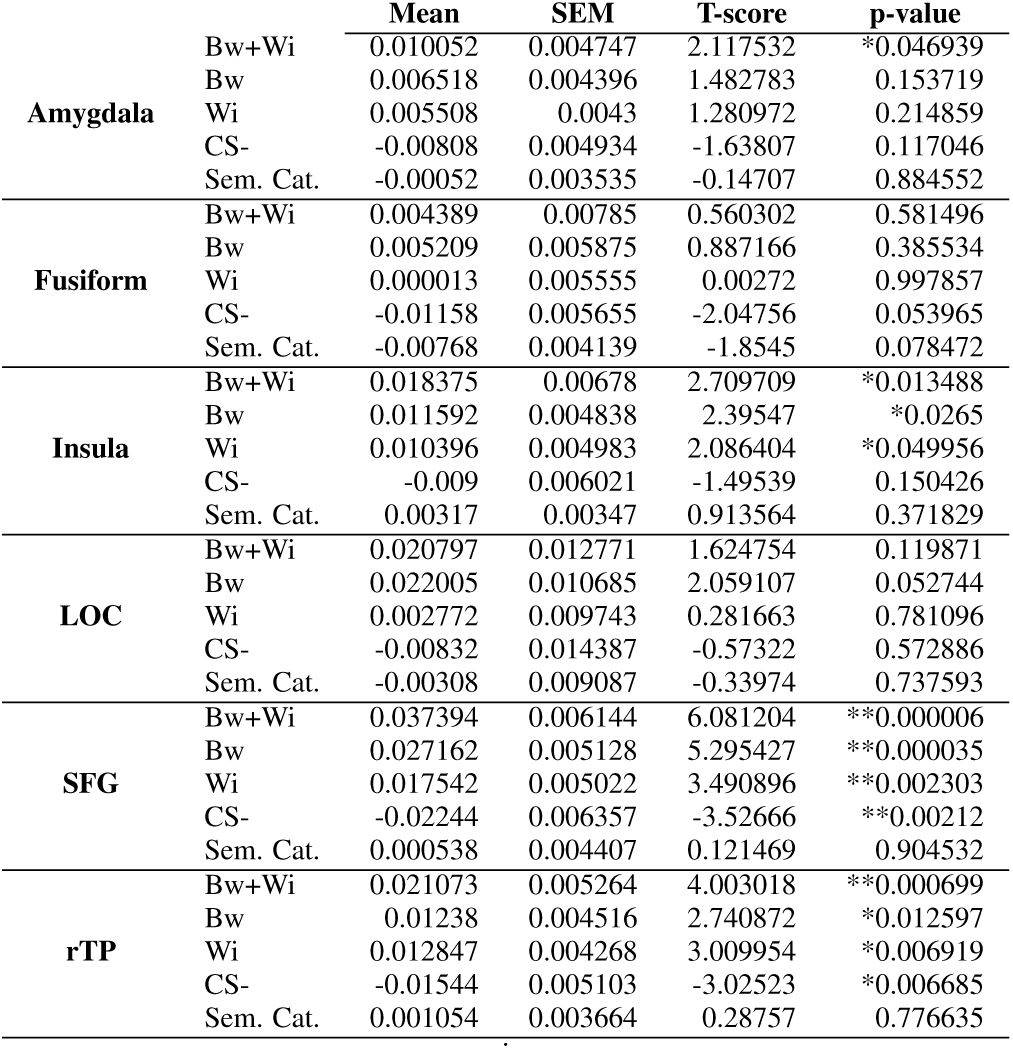
Descriptive statistics of the contrast Conditioned > Baseline for the five models at all six regions. *p_unc_< 0.05, **p_FDR_< 0.05

Additionally, following fear-conditioning, the Bw+Wi model outperformed the Bw model (which was the next best model) in the SFG (t_20_ = 2.34, p_unc_ = 0.0296) but not in the rTP (t_20_ = 1.198, p_unc_ = 0.1225). However, in both the SFG and the rTP, the Bw+Wi model yielded a greater change from the baseline phase to the conditioned phase as compared to the Bw model (t_20_ = 2.43, p_unc_ = 0.0245; t_20_ = 2.42, p_unc_ = 0.0253, respectively).

### Lateral occipital cortex prefers the semantic model, regardless of experimental phase

Given previously demonstrated pattern similarity changes in object-selective cortex, we wondered whether such regions would also yield similar results with respect to between-category similarity changes. However, the LOC showed a preference for the semantic model, which consistently outperformed the next-most-preferred model (the Wi model, which is a similar model [Spearman’s *ρ* ≈ 0.76]) at all three experimental phases (Bl: t_20_ = 2.97, p_unc_ = 0.0075; Co: t_20_ = 2.64, p_unc_ = 0.016; Ex: t_20_ = 4.85, p_unc_ = 9.66 × 10^−5^). Unlike the SFG and ATL, there was not sufficient evidence to demonstrate that the Bw+Wi, the Bw, or the Wi models were modulated by fear-conditioning (t_20_ = 1.62, p = 0.120; t_20_ = 2.06, p = 0.053; t_20_ = 0.282, p = 0.781, respectively) in the LOC. Despite the trend of the Bw model from the baseline phase to the conditioned phase, it is interesting to note that the Bw model tended to anticorrelate with activity patterns in the LOC, predominantly during the baseline and extinction phases (t_20_ = −2.94, p_unc_ = 0.008; t_20_ = −2.68, p_unc_ = 0.015, respectively), but less so at the conditioned phase (t_20_ = −1.87, p_unc_ = 0.076).

### No evidence for relationship between anxiety scores and conditioning effects

Using the STAI data that the participants provided, we Pearson correlated the STAI scores with the changes in correlation, between baseline and conditioned, for the Bw+Wi, Bw, Wi, and CS-models in the SFG and the Bw+Wi model in the rTP (i.e., the models that survived the FDR-correction). Ultimately, we were unable to find such relationships between these variables, with the exception of a possible correlation between the Bw model and the SFG’s correlation changes (r = 0.43, p = 0.053). See Table 3 for a full list of the correlation results.

**Table 3.** Results from Pearson correlating the STAI scores with the changes in the FDR-corrected models’ explanatory value. P-values are in parentheses.

|  | SFG |  | rTP |  |
| --- | --- | --- | --- | --- |
|  | STAI-S | STAI-T | STAI-S | STAI-T |
| Bw+Wi | 0.113 (0.627) | -0.041 (0.860) | - | - |
| Bw | 0.428 (0.053) | -0.217 (0.344) | 0.357 (0.112) | -0.003 (0.990) |
| Wi | 0.277 (0.225) | -0.285 (0.211) | - | - |
| CS- | 0.263 (0.250) | -0.021 (0.930) | - | - |

## Discussion

This study aimed to investigate how information represented in the human brain is altered as a function of aversively conditioning exemplars from multiple semantic categories. Prior work combining fMRI and fear-conditioning has often focused on differences between a CS+ and CS-, either in terms of category representations [10] or generalization gradients [9, 28]. Conducting an experiment with three semantic categories (2 CS+’s and 1 CS-), we employed the framework of representational similarity analysis (RSA) [11] to test whether fear-conditioning would also generalize *between* categories and whether such between-category effects would also be present in the same brain regions in which within-category effects have been reported.

Our primary findings revealed that activity patterns within the SFG and the rTP—two regions previously reported to have shown pattern changes following aversive-learning [8, 24])—were best explained by the Bw+Wi model, indicating increased similarity of both between- and within-category exemplars after fear-conditioning. Such findings suggest that fear-generalization, as measured by the similarity of activity patterns, can extend beyond the level of single categories in certain brain regions, and that regions like the SFG and rTP play a more general role in processing fear information (as our results imply an additional supracategorical effect in that not all participants were conditioned to the same two categories). This role may be intrinsically linked to semantic cognition, given that previous studies have related semantic processing to functional and structural properties of the anterior temporal lobes [29–31] as well as to multivoxel activity patterns in the SFG [32, 33]. Alternatively, these regions’ role in fear-generalization may underlie affective processes, as diverse aspects of emotion information processing have been linked to both the SFG [34–37] and the anterior temporal lobe [38–40]. This second point is bolstered by the fact that the results we reported within the insula reflected those from the SFG and the rTP. Although our insular results did not surpass the FDR-corrected statistical thresholds, numerous studies have linked the insula to affective processing [41–47].

Activity patterns within the object-selective LOC were best explained by the semantic category model, regardless of the experimental phase. This observation of category-level representations within the LOC fits with prior literature [48, 49] and is also further supported by the finding that the Bw model (the only model containing no within-category similarity) was generally anticorrelated with LOC activity patterns (though this particular point results from uncorrected p-values, so we refrain from drawing conclusions based solely on these statistics). Interestingly, we did not find that activity patterns within object-selective cortex or the amygdala were modulated by fear-conditioning, which stands in contrast to previous findings [10]. However, we would not call this result a failure to replicate prior work because it is, in fact, unclear to what degree our results are directly comparable to those of previous reports, as the experimental paradigms differed. Specifically, our experiment resulted in two of three categories being aversively conditioned; thus, the *majority* of stimulus material would yield a higher shock expectancy, as opposed to one-half of stimulus material in studies that contain one CS+ category and one CS-category.

This aspect of the design brings up two points: first, the representations within the SFG and rTP may already abstract away from semantic categories altogether and instead reflect fear information. This idea is supported by the fact that we fear-conditioned two distinct semantic categories. As such, any brain regions that appear to reflect some form of between-category conditioning may be reflecting supracategorical or more generic information pertaining to the affective processes underlying the fear circuitry. Additionally, given that different participants were conditioned to different sets of categories, the group-level results we present add further evidence to this point. This interpretation is also in line with the previous finding of increased similarity between a semantically unrelated CS+ category and a phobic category in the rTP [8].

Second, investigations that employ more fear-conditioned than neutral categories may be studying processes in a fundamentally different state than those experiments employing an equal number of fear-conditioned and neutral categories. This potentially distinction may result from participants internally normalizing the aversive nature of the “environment”, thereby increasing the saliency or the value of the *safe* category at some cognitive level. In such cases, the information entropy [50] is lower, giving rise to potentially different physiological processing of threat-related (un)certainty [51–54].

In summary, our experiment revealed increases in both between-and within-category similarity in more anterior regions of the brain: specifically, the superior frontal gyrus and the right temporal pole. Both of these areas have been previously implicated in diverse aspects of emotion information processing; as such, it is conceivable that these regions play a role in representing abstract information pertinent to fear-learning. This notion places a spotlight on such regions when translating related experimental designs to the clinical domain in order to explore potential functional imaging-related biomarkers of anxiety disorders.

## Acknowledgments

The authors would like to thank Dasa Zeithamova-Demircan for contributing stimulus material. The formatting style for this manuscript derives from the “HenriquesLab bioRxiv template” by Ricardo Henriques, used under CC BY 4.0.

## Author contributions

SL, MK, and JS designed the study. SL and MK acquired data, analyzed data, and drafted the manuscript. SL, MK, and JS revised the manuscript. RR contributed experiment resources. All authors approved the final version of the manuscript.

## Data availability

The data that support the findings of this study are available from the corresponding author upon reasonable request.

